# Postural control in girls practicing volleyball is different than in non-playing peers: comparison of data on the center of pressure and ground reaction forces

**DOI:** 10.1101/2023.07.28.550870

**Authors:** Dorota Borzucka, Krzysztof Kręcisz, Michał Kuczyński

## Abstract

A comprehensive explanation of the relationship between postural control and athletic performance requires evaluating body balance in athletes of different performance levels. To fill this gap in relation to volleyball, the aim of this study was to compare the balance of intermediate adolescent female players (VOL, n=61) with inactive peers (CON, n=57). The participants were investigated in normal quiet stance during 30 s trials on a Kistler force plate. The traditional spatial (amplitude and mean speed) and temporal (frequency, fractality and entropy) indices were computed for ground reaction forces (GRF) and center-of-pressure (COP) time-series. The spatial parameters of the both time-series did not discriminate the two groups. However, the temporal GRF parameters revealed much lower values in VOL than in CON (p<.0001). This leads to three important conclusions regarding posturography applications. First, GRF and COP provide different information regarding postural control. Second, measures based on GRF are more sensitive to changes in balance (at least those related to volleyball training and perhaps to similar training and sports activity regimens). And third, the indicators calculated on the basis of these two time series can complement each other and thus enrich the insight into the relationship between balance and sports performance level.

## INTRODUCTION

Volleyball is a team game that puts high demands on players in terms of motor skills. The actions take place very quickly, there is no way to hold the ball, so the player has fractions of a second to make a decision (Borzucka et al., 2020a, 2020b). It would seem that the ability to control balance is not particularly important in volleyball. Nothing could be more wrong. Skillful control of postural stability is the basis of many activities (Agostini et al., 2013). Players have to hit the ball in different positions, both in stable and disturbed balance (Kuczyński et al., 2009). Although most points are scored in situations where there is no contact with the ground (attack, block, jump serve), success is often determined by less spectacular plays, such as receiving a serve or defense (Borzucka et al., 2020a, 2020b). In these activities, a stable attitude is always sought (Agostini et al., 2013; Kuczyński et al., 2009). This is to ensure greater precision of the performance of a given element. It is not surprising, therefore, that trainers pay more and more attention to balance training. It aims not only to improve the performance of the team, but also to prevent injuries, which is confirmed by studies conducted among athletes of various sports disciplines (Agostini et al., 2013; McGuine & Keene, 2006; Petersen & Hölmich, 2005; Söderman et al., 2000; Verhagen et al., 2004).

Studying postural control in athletes can provide insight into the development of specific postural strategies that are particularly relevant in a given sport (Agostini et al., 2013; Malliou et al., 2010; Sannicandro et al., 2014). The aim of our research so far was to investigate postural strategies of volleyball players at various levels of sports advancement. We surveyed volleyball players of the 2nd league (Kuczyński et al., 2009), as well as volleyball players representing the championship level (Polish women’s and men’s national team) (Borzucka et al., 2020a, 2020b).

Some of the desired features and postural skills have been confirmed, e.g. higher automaticity, better use of sensory signals, adaptability to changing environment and high exploratory capabilities (Bieć & Kuczyński, 2010; Piątek-Krzywicka et al., 2022; Sipko & Kuczyński, 2013). Yet, it is not a completely consistent picture and some doubts still arise. Importantly, however, very large differences in the sports level of these previous groups revealed the possibility of transient phases in the development of their postural strategies. For example, players from the second league did not have a significant advantage over physically active peers. Although the former ones had a small sway amplitude, this was at the expense of higher sway speed (Kuczyński et al., 2009). This could indicate the stage of building new postural strategies towards, for example, optimizing the compromise between the accuracy and the speed of using afferent signals.

To confirm this assumption, it is worth analyzing the changes in postural control that appear at the early stage of the athlete’s career. The possible direction of changes may be the best visible here, and moreover, it may determine the subsequent development of postural control in order to equip it with elements supporting the effectiveness of playing volleyball. Thus, we considered it important to investigate postural control in young girls who already have some experience in this sport.

Previous studies have been conducted solely on the basis of the center-of-pressure (COP) data. However, the results of our own observations and those of other authors indicate that the discriminating power of this computational approach is insufficient (Caballero et al., 2021; Chapman et al., 2008; Paillard et al., 2002). Quite often the results collected in quiet normal stance while comparing groups with very different sports levels (i.e. with hypothetically significant differences in balance) are ambiguous or, at most, wind up with insignificant differences and small effect sizes (Andreeva et al., 2020; Kiers et al., 2013). One way to improve this situation is to test subjects in more demanding experimental conditions (Srihi et al., 2022; Zemková & Kováčiková, 2022) and preferably in those that are similar to tasks performed during sports competitions or training. Another approach could be to use new indicators that are known to be strongly associated with postural control.

Following the second option, we decided to include the horizontal ground reaction forces (HGRF) in addition to using the COP to assess balance. Goldie et al., (Goldie et al., 1989) were the first to publish similar results and reported that the parameters calculated from the HGRF have better test-retest reliability and sensitivity to changes in quiet stance than COP parameters. Unfortunately, this topic has been scarcely addressed by other researchers except Karlsson and Frykberg (Karlsson & Frykberg, 2000) and Onell (Onell, 2000), and more recently Minamisawa et al. (Minamisawa et al., 2012) who confirmed the specific and additional value of these measures compared to the COP parameters. Nevertheless, to this day it is not clear how these measures work, what they help to detect in the field of postural control and what is their relationship with the COP parameters. For more information on this issue, we recalculated the source data from some of our previously published articles. The results confirmed our assumptions that the HGRF indices are in no way inferior to the COP indices. In addition, the results obtained by both methods may differ significantly, even when it comes to the direction of changes in the same parameters of both time-series under the influence of experimental manipulations. This confirmed our belief that the inclusion of the HGFR indices in the calculations should result in additional information about the postural control of the subjects.

Therefore, the aim of this study was to compare postural control in young female volleyball players with their non-training counterparts using the COP and HGRF time-series as data to calculate indices of their spatial and temporal structure. We hypothesized that postural control would be different in both groups, and any differences should result from the specific requirements of playing volleyball. We also hypothesized that the HGRF indices would better differentiate the two groups than the corresponding COP indices.

## MATERIALS AND METHODS

### Participants

The study included 61 young female volleyball players (VOL): age 12.8 ± 1.4 yr; height 162.5 ± 9.5 cm; weight 52.8 ± 11.4 kg. The average training period was ?? ± ??yr. Trainings were held 3-4 times a week, in addition to matches and tournaments that were played. The control group consisted of 57 girls (CON): age 12.9 ± 1.5 yr; height 156.1 ± 8.7 cm; weight, 47.9 ± 16.2 kg, who did not undertake systematic physical activity (except for mandatory physical education classes at school). All respondents described their health as good or very good.

This study was approved by the Bioethics Committee of Opole Chamber of Physicians in Opole. Written informed consent was obtained from all participants and their parents or legal guardian(s) prior to participation. The objectives, procedures, and methods were explained in full and the subjects were informed that they could withdraw at any time. All methods were carried out according to the relevant guidelines and regulations.

### Methods

Postural control was assessed on a Kistler force platform (Kistler 9286AA, Winterthur, Switzerland). Ground reaction forces were recorded, and the coordinates of the COP data were calculated for 20 s, at a sampling frequency of 100 Hz separately for frontal (ML) and sagittal (AP) planes. The study protocol was the same as in (Borzucka et al., 2020b).

Each participant performed one trial of standing quietly on a hard surface in an upright position in a neutral and comfortable posture with arms relaxed on the sides and eyes open. They were instructed to stand as still as possible for 20 seconds. The position of the feet (5 cm apart) was standardized on the surface to ensure reproducibility among the participants. The data acquisition started when the participant indicated that he was ready.

### Parameters

Based on HGRF and COP recordings, postural control parameters were calculated. These were (Duarte & Freitas, 2010): standard deviation (SD), mean speed (MV), sample entropy (SE), frequency (FRE = MV/(2*π*SD)). Parameters for estimating sample entropy were based on the median value of the relative error that resulted in the selection of pattern length of m= 3 and error tolerance of r= 0.02 for COP and m=3 and r=0.15 as optimal parameters for both the ML and AP planes (normalized to unit variance) of all subjects and tasks (Bieć et al., 2014; Lake et al., 2002).

### Statistics

All dependent variables were subjected to T-Tests. If the assumption of equal variance was not met Welch’s test was used to compare groups. Similarly, if the normality assumption was not met (assessed graphically through Q-Q and residual histogram plots) Mann–Whitney U group independent test was used. Cohen’s effect sizes were also reported. The level of significance was set at *p* < 0.05. The analyses were performed using the jamovi (ver. 2.3.24) software.

## RESULTS

The results are presented in Table 1. The COP parameters found only one difference between groups, namely in the ML mean sway speed which was higher in VOL than in CON with moderate effect size. In contrast, the HGRF parameters revealed many intergroup differences, which always occurred in both planes. In particular, large effect sizes were observed for sway entropy and frequency indicating smaller values in VOL, while moderate effect sizes appeared for sway amplitude (higher values in VOL).

**Table 1.** Parameters of the horizontal forces (HGRF) and COP in the frontal (ML) and sagittal (AP) direction for two groups (volleyball and control) in quiet bipedal stance. The entries are mean values (SD).

| HGRF |  |  |  | COP |  |  |  |
| --- | --- | --- | --- | --- | --- | --- | --- |
| ML |  | AP |  | ML |  | AP |  |
| Volleyball | Control | Volleyball | Control | Volleyball | Control | Volleyball | Control |
| AMPLITUDE (N) |  |  |  | AMPLITUDE (mm) |  |  |  |
| 0.58 (0.23) | 0.46 (0.19) | 0.72 (0.21) | 0.59 (0.20) | 3.02 (1.29) | 2.81 (1.62) | 4.35 (1.70) | 3.81 (1.32) |
| p=0.0023/-0.57 |  | p=0.0011/-0.62 |  |  |  |  |  |
| MEAN SPEED (N/s) |  |  |  | MEAN SPEED (mm/s) |  |  |  |
| 7.82 (1.56) | 8.33 (2.12) | 8.87 (2.49) | 9.21 (3.16) | 9.27 (2.49) | 7.89 (2.44) | 12.77 (2.60) | 11.94 (3.98) |
|  |  |  |  | p=0.0068/0.56 |  |  |  |
| ENTROPY (-) |  |  |  | ENTROPY (-) |  |  |  |
| 0.81 (0.24) | 1.03 (0.30) | 0.65 (0.14) | 0.85 (0.18) | 1.12 (0.30) | 1.01 (0.37) | 1.15 (0.35) | 1.16 (0.35) |
| p<0.0001/0.81 |  | p<0.0001/1.21 |  |  |  |  |  |
| FREQUENCY (Hz) |  |  |  | FREQUENCY (Hz) |  |  |  |
| 2.39 (0.73) | 3.29 (1.30) | 2.05 (0.55) | 2.57 (0.65) | 0.54 (0.16) | 0.52 (0.20) | 0.52 (0.19) | 0.53 (0.21) |
| p<0.00001/0.86 |  | p<0.00001/0.88 |  |  |  |  |  |

The p-values are for the inter-group comparisons; the number after the slash indicates Cohen's effect size

Examples of HGRF and COP time series are shown in Figure 1 that illustrate differences in the amplitude and regularity of the signals.

**Figure 1.**
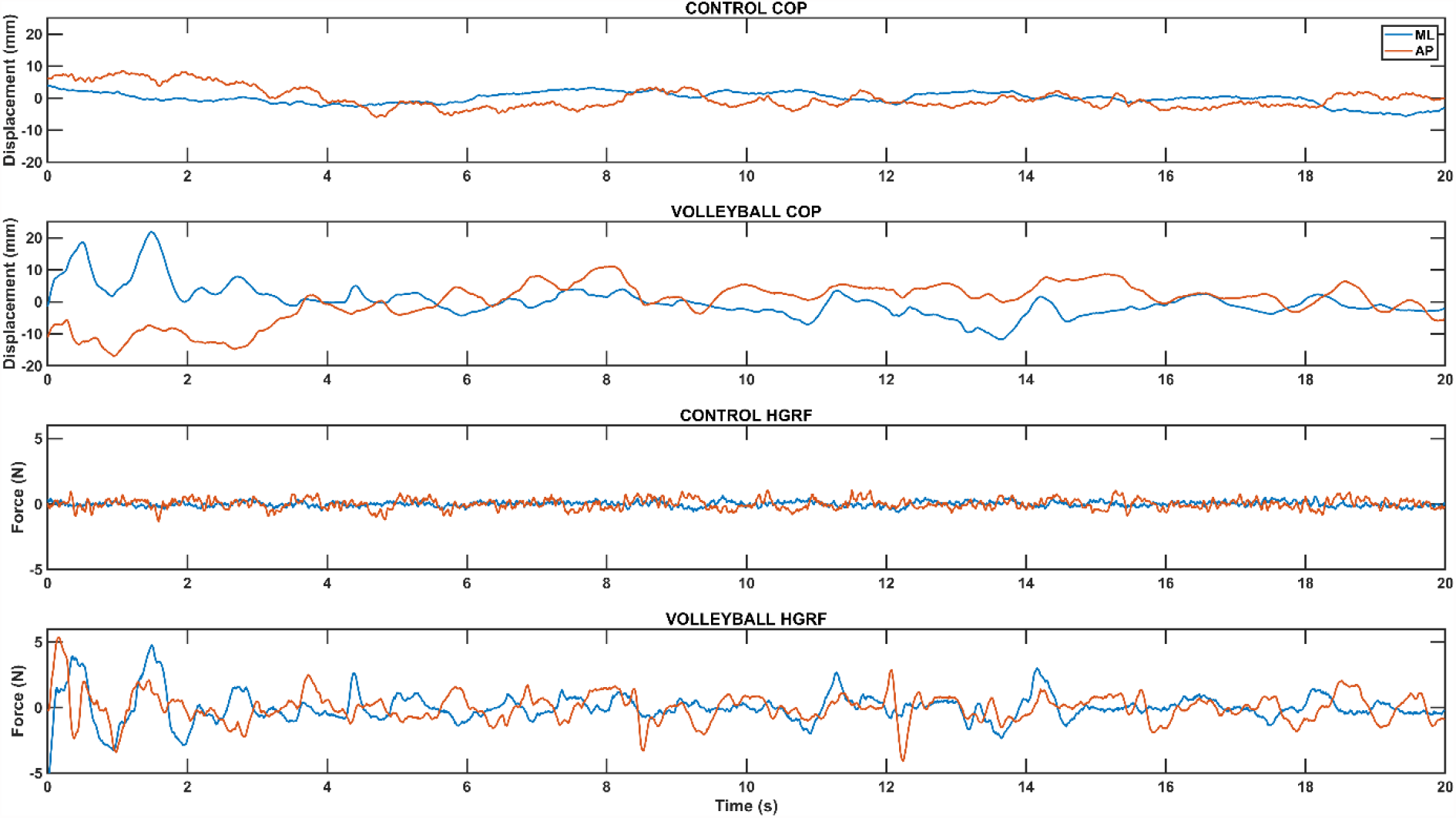
Selected time series of HGRF and COP signals from two study participants showing differences in dynamic structure during quiet bipedal stance. ML – frontal plane, AP – sagittal plane, CONTROL COP – COP time series of participant from control group, VOLLEYBALL COP – COP time series of participant from volleyball group, CONTROL HGRF – the horizontal force time series of participant from control group, VOLLEYBALL HGRF – the horizontal force time series of participant from volleyball group. The mean waveform has been subtracted from the individual waveforms.

## DISCUSSION

The purpose of this study was to compare postural control in bipedal stance between adolescent female volleyball players and their non-training peers. Being aware of the insufficient discriminating power presented by the COP measures collected during typical everyday postural tasks such as standing still on both legs, we additionally used the same measures calculated directly from the horizontal ground reaction forces time-series. As expected, only one COP-based index, namely COP speed in the ML axis, showed moderate differences between the two groups. However, the direction of this difference was opposite to the expected one, i.e. VOL had higher speed than CON. On the other hand, the HGRF calculations revealed substantial intergroup differences for all indices except for speed, which values were similar in both groups. This new information is very interesting in itself and may help shed light on the processes involved in the development of postural control in young female volleyball players.

As can be seen, neither enriching the COP parameter set with frequency and sample entropy measures nor additional simple calculations on the HGRF help in understanding the detected differences between the groups. It is therefore worth taking advantage of the additional results based on entropy and frequency of the HGRF.

Consider first the entropy, which was rather unexpectedly lower in VOL, i.e. showed a less irregular temporal structure of the HGRF time-series than in CON. Most studies using this measure to assess COP dynamics claim that increased, not decreased, irregularity indicates beneficial changes in postural control (Akbaş et al., 2022; Piątek-Krzywicka et al., 2022; Terada et al., 2018). However, there is no shortage of authors who recognize the inverse relationship, i.e. indicate improved (or better) stability at lower entropy values (Hadad et al., 2020; Raffalt et al., 2021). Thus, this problem has not yet been solved for the COP, let alone for the HGRF, for which no similar comparisons have been made at all. It seems that the best way to make such comparisons will be to base them on the specificity of postural behavior of volleyball players and the challenges of this sport for body stability.

An argument in favor of lower entropy in VOL may be the specific attitude of volleyball players on the court with their knees bent and slightly leaning forward, which is a preparation for the next action whatever it might be. A similar strategy can be observed while standing on a foam surface (Piątek-Krzywicka et al., 2022; Sipko & Kuczyński, 2013), which serves to better exploit the afferent signals from the feet and reflect a purposeful attempt to increase the amount feedback available (Strang et al., 2011). These attempts are accompanied by conscious, ample and systematic swaying of the body with fairly constant parameters. The superposition of such a more predictable signal with normal chaotic-like involuntary swaying must lead to greater regularity in the resultant sway, i.e. to lower COP entropy. Thus, there is at least one reason for a decreased COP entropy in the training group: specific requirements of volleyball game. Following the example of a few authors (e.g. (Raffalt et al., 2021)), we will refrain here from indicating this “better” entropy. They are simply different in the two groups, which is a satisfactory result in view of the purpose of this study.

It is very often argued that higher entropy is a sign of greater automaticity of postural control (with less attention) and therefore should be equated to adopting better postural strategies (Piątek-Krzywicka et al., 2022; Richer & Lajoie, 2020; Szafraniec et al., 2018). Does better automaticity really mean better control? We believe it depends on the situation, so sometimes it is useful and other times it can be a hindrance. Firstly, such conclusions result from specific experiments concerning manipulation of attention resources, while our descriptive study did not introduce such manipulations and nevertheless revealed much lower entropy in the VOL group. Perhaps interpretations based on a combination of automaticity and entropy are only a small part of much more complex relationships. For example, it is known that easier postural tasks may exhibit lower COP entropy (Lubetzky et al., 2015; Saraiva et al., 2023). In a similar vein, lower entropy was found in young than in elderly subjects (Borg & Laxåback, 2010) and in non-fallers than in fallers (Montesinos et al., 2018). In this context, the lower entropy of our female volleyball players would mean that quiet stance is easier for female volleyball players than for CON and/or VOL perform this task with greater confidence. And this easiness does not have to be in any way related to the level of automatic control. It may be the result of some other beneficial adaptations.

Secondly, how would a higher degree of postural control automaticity support our VOL during a match? Even if some attentional resources are released through increased automaticity, the need for constant vigilance and supervision (during a game) will likely set them back. Further, control of automatic processes is difficult and requires substantial effort (Moors & De Houwer, 2006). It follows that while automaticity may be convenient and useful in several everyday situations, this asset is questionable on a volleyball court. And even more - it can be a serious obstacle in the conditions of fierce competition. Thus, lower sway entropy (lower irregularity) may indicate an advantage in VOL’s postural control over CON, especially in the face of stability challenges during a match. This is notably true of the trade-off between stability and maneuverability (Acasio et al., 2017; Borzucka et al., 2020b), which is particularly desirable and at the same time difficult to achieve during competitions.

The COP frequency exhibited no difference between the two groups in this study. In contrast, the frequency of HGRF was much lower in VOL than in CON, which suggests important changes in VOL under the influence of sport. We recently compared top-level female volleyball players with non-training peers (Borzucka et al., 2020a) and also found no differences between sway frequencies while the COP-COM frequency was larger in athletic group. It is important to note that COP-COM is proportional to horizontal accelerations (Masani et al., 2007; Winter et al., 1998) which can be derived from the HGRF and are directly associated with the neural input to the postural control system. Thus, the calculations based on the HGRF correspond to the respective COP-COM calculations. In another work (Borzucka et al., 2020b), we reported much higher COP frequency in top-level male volleyball players as compared with non-athletes which was interpreted as means to alleviate postural sway. This was accompanied by a decrease in the AP COP-COM frequency in athletes. Lastly, we compared second league volleyball players with non-active controls and found higher AP COP-COM frequency in athletes (Kuczyński et al., 2009). It follows quite clearly that it is impossible to capture any constant pattern of frequency changes as a function of the sport performance level, at least in volleyball.

To solve this conundrum, we proposed speculatively that any unexpected changes in FRE may be tentatively treated as a sign of transient changes in the choice of postural strategies (Borzucka et al., 2020a). This idea does not seem to be far from reality, because there are many factors that can affect or provoke changes in postural control at any level of experience. However, scientific support would be needed to confirm such speculation. Some papers linking motor variability with motor learning come to our aid here. For instance, recent findings show that increased variability improves learning and supports exploration as a tool to search for new strategies (Abram et al., 2022; Roemmich & Bastian, 2022). Additionally, it has been reported that specific changes in the frequency distribution of a neural drive can modulate motor variability associated with that drive (Park et al., 2017) and that forward leaning posture relates to low-frequency neural inputs (Watanabe et al., 2019).

Considering these last suggestions, let us recall the relevant results (HGRF) of this study. Horizontal forces showed greater variability in VOL, and the force frequency measures decreased in this group. The first observation should be interpreted as a manifestation of information seeking for the purpose of further exploration in the space of possible solutions, which is characteristic of the process of learning and adaptation (Abram et al., 2022). On the other hand, the second observation proves the increased importance of lower FRE in the HGRF spectrum. This is not entirely unambiguous, because it could be caused by both an increase in the amplitude of lower frequencies as well as a decrease in higher ones. Putting these details aside for now, this means, first, a voluntary increase in horizontal forces that has been directly measured. Secondly, it indicates a tendency to rock regularly in a slightly forward-leaning position. And lastly, this is in line with Richer and Lajoie (Richer & Lajoie, 2020) who interpreted the lower frequency as a sign of decreased automaticity, which in turn supports our findings of reduced entropy.Summing up, we believe that we have drawn reasonable inferences of the contrast between postural control in VOL and CON. It is worth emphasizing that these results were obtained only on the basis of HGRF analysis using indices commonly used for COP.

There are two major limitations in this study that could be addressed in future research. First, in trying to understand the reasons for the different dynamics (temporal structure) of the HGRF in the two groups, we drew to some extent from previous interpretations of similar changes observed in the COP. Therefore, it is important to be aware that these interpretations are not necessarily correct because the HGRF contains more information about postural control than the COP (the frequency band of the first signal is about 4 times larger than the second). This surplus of information, the meaning of which is yet unknown, may modify our (still incomplete) picture of the relationship between experimental conditions and entropy. On the other hand, it can also contribute to a better understanding of non-linear measures of COP. The second limitation concerns the central tendency measure for the HGRF frequency. While various such point measures exist and are useful in many applications, their use in the study of human neurophysiology appears to be limited and even confusing. In order to avoid undesirable ambiguities, it is recommended to divide the entire frequency band into several parts. This will allow, to some extent, to isolate changes in frequency depending on the neural control strategy used by subjects during quiet stance.

## CONCLUSION

The frequency and entropy of the HGRF time-series were very effective in distinguishing the volleyball players from the control group, which was completely unsuccessful when analyzing the COP only. This means that despite very similar COP indices in the two compared groups, the processes at the interface of neural control of balance and sway may differ significantly. Most likely, it accounts for a different way of processing information and/or different control modes convenient for each group. The HGRF parameters show greater sensitivity to possible differences in postural control and may be particularly useful in tracking changes in the maintenance of upright stance over short periods of time.

## Notes

### Competing Interest Statement

The authors have declared no competing interest.

## REFERENCES

Abram, S. J., Poggensee, K. L., Sánchez, N., Simha, S. N., Finley, J. M., Collins, S. H., & Donelan, J. M. (2022). General variability leads to specific adaptation toward optimal movement policies. Current Biology, 32(10), 2222–2232.e5. https://doi.org/10.1016/j.cub.2022.04.015

Acasio, J., Wu, M., Fey, N. P., & Gordon, K. E. (2017). Stability-maneuverability trade-offs during lateral steps. Gait & Posture, 52, 171–177. https://doi.org/10.1016/j.gaitpost.2016.11.034

Agostini, V., Chiaramello, E., Canavese, L., Bredariol, C., & Knaflitz, M. (2013). Postural sway in volleyball players. Human Movement Science, 32(3), 445–456. https://doi.org/10.1016/j.humov.2013.01.002

Akbaş, A., Marszałek, W., Drozd, S., Czarny, W., Król, P., Warchoł, K., Słomka, K. J., & Rzepko, M. (2022). The effect of expertise on postural control in elite sport ju-jitsu athletes. BMC Sports Science, Medicine and Rehabilitation, 14(1), 86. https://doi.org/10.1186/s13102-022-00477-3

Andreeva, A., Melnikov, A., Skvortsov, D., Akhmerova, K., Vavaev, A., Golov, A., Draugelite, V., Nikolaev, R., Chechelnickaia, S., Zhuk, D., Bayerbakh, A., Nikulin, V., & Zemková, E. (2020). Postural Stability in Athletes: The Role of Age, Sex, Performance Level, and Athlete Shoe Features. Sports (Basel, Switzerland), 8(6). https://doi.org/10.3390/sports8060089

Bieć, E., & Kuczyński, M. (2010). Postural control in 13-year-old soccer players. European Journal of Applied Physiology, 110(4), 703–708. https://doi.org/10.1007/s00421-010-1551-2

Bieć, E., Zima, J., Wójtowicz, D., Wojciechowska-Maszkowska, B., Krȩcisz, K., & Kuczyński, M. (2014). Postural stability in young adults with down syndrome in challenging conditions. PLoS ONE, 9(4), e94247. https://doi.org/10.1371/journal.pone.0094247

Borg, F. G., & Laxåback, G. (2010). Entropy of balance--some recent results. Journal of Neuroengineering and Rehabilitation, 7, 38. https://doi.org/10.1186/1743-0003-7-38

Borzucka, D., Kręcisz, K., Rektor, Z., & Kuczyński, M. (2020a). Postural control in top-level female volleyball players. BMC Sports Science, Medicine & Rehabilitation, 12, 65. https://doi.org/10.1186/s13102-020-00213-9

Borzucka, D., Kręcisz, K., Rektor, Z., & Kuczyński, M. (2020b). Differences in static postural control between top level male volleyball players and non-athletes. Scientific Reports, 10(1), 19334. https://doi.org/10.1038/s41598-020-76390-x

Caballero, C., Barbado, D., Hérnandez-Davó, H., Hernández-Davó, J. L., & Moreno, F. J. (2021). Balance dynamics are related to age and levels of expertise. Application in young and adult tennis players. PloS One, 16(4), e0249941. https://doi.org/10.1371/journal.pone.0249941

Chapman, D. W., Needham, K. J., Allison, G. T., Lay, B., & Edwards, D. J. (2008). Effects of experience in a dynamic environment on postural control. British Journal of Sports Medicine, 42(1), 16–21. https://doi.org/10.1136/bjsm.2006.033688

Duarte, M., & Freitas, S. M. S. F. (2010). Revision of posturography based on force plate for balance evaluation. Revista Brasileira de Fisioterapia (Sao Carlos (Sao Paulo, Brazil)), 14(3), 183–192. http://www.ncbi.nlm.nih.gov/pubmed/20730361

Goldie, P. A., Bach, T. M., & Evans, O. M. (1989). Force platform measures for evaluating postural control: reliability and validity. Archives of Physical Medicine and Rehabilitation, 70(7), 510–517. http://www.ncbi.nlm.nih.gov/pubmed/2742465

Hadad, A., Ganz, N., Intrator, N., Maimon, N., Molcho, L., & Hausdorff, J. M. (2020). Postural control in karate practitioners: Does practice make perfect? Gait and Posture, 77, 218–224. https://doi.org/10.1016/j.gaitpost.2020.01.030

Karlsson, A., & Frykberg, G. (2000). Correlations between force plate measures for assessment of balance. Clinical Biomechanics (Bristol, Avon), 15(5), 365–369. https://doi.org/10.1016/s0268-0033(99)00096-0

Kiers, H., van Dieën, J., Dekkers, H., Wittink, H., & Vanhees, L. (2013). A systematic review of the relationship between physical activities in sports or daily life and postural sway in upright stance. Sports Medicine (Auckland, N.Z.), 43(11), 1171–1189. https://doi.org/10.1007/s40279-013-0082-5

Kuczyński, M., Rektor, Z., & Borzucka, D. (2009). Postural Control in Quiet Stance in the Second League Male Volleyball Players. Human Movement, 10(1). https://doi.org/10.2478/v10038-008-0025-4

Lake, D. E., Richman, J. S., Pamela Griffin, M., & Randall Moorman, J. (2002). Sample entropy analysis of neonatal heart rate variability. American Journal of Physiology - Regulatory Integrative and Comparative Physiology, 283(3 52-3), R789–R797. https://doi.org/10.1152/ajpregu.00069.2002

Lubetzky, A. V, Price, R., Ciol, M. A., Kelly, V. E., & McCoy, S. W. (2015). Relationship of multiscale entropy to task difficulty and sway velocity in healthy young adults. Somatosensory and Motor Research, 32(4), 211–218. https://doi.org/10.3109/08990220.2015.1074565

Malliou, V. J., Beneka, A. G., Gioftsidou, A. F., Malliou, P. K., Kallistratos, E., Pafis, G. K., Katsikas, C. A., & Douvis, S. (2010). Young tennis players and balance performance. Journal of Strength and Conditioning Research, 24(2), 389–393. https://doi.org/10.1519/JSC.0b013e3181c068f0

Masani, K., Vette, A. H., Kouzaki, M., Kanehisa, H., Fukunaga, T., & Popovic, M. R. (2007). Larger center of pressure minus center of gravity in the elderly induces larger body acceleration during quiet standing. Neuroscience Letters, 422(3), 202–206. https://doi.org/10.1016/j.neulet.2007.06.019

McGuine, T. A., & Keene, J. S. (2006). The effect of a balance training program on the risk of ankle sprains in high school athletes. The American Journal of Sports Medicine, 34(7), 1103–1111. https://doi.org/10.1177/0363546505284191

Minamisawa, T., Sawahata, H., Takakura, K., & Yamaguchi, T. (2012). Characteristics of temporal fluctuation of the vertical ground reaction force during quiet stance in Parkinson’s disease. Gait & Posture, 35(2), 308–311. https://doi.org/10.1016/j.gaitpost.2011.09.106

Montesinos, L., Castaldo, R., & Pecchia, L. (2018). On the use of approximate entropy and sample entropy with centre of pressure time-series. Journal of NeuroEngineering and Rehabilitation, 15(1), 116. https://doi.org/10.1186/s12984-018-0465-9

Moors, A., & De Houwer, J. (2006). Automaticity: A theoretical and conceptual analysis. Psychological Bulletin, 132(2), 297–326. https://doi.org/10.1037/0033-2909.132.2.297

Onell, A. (2000). The vertical ground reaction force for analysis of balance? Gait & Posture, 12(1), 7–13. https://doi.org/10.1016/s0966-6362(00)00053-9

Paillard, T., Costes-Salon, C., Lafont, C., & Dupui, P. (2002). Are there differences in postural regulation according to the level of competition in judoists? British Journal of Sports Medicine, 36(4), 304–305. https://doi.org/10.1136/bjsm.36.4.304

Park, S. H., Casamento-Moran, A., Yacoubi, B., & Christou, E. A. (2017). Voluntary reduction of force variability via modulation of low-frequency oscillations. Experimental Brain Research, 235(9), 2717–2727. https://doi.org/10.1007/s00221-017-5005-5

Petersen, J., & Hölmich, P. (2005). Evidence based prevention of hamstring injuries in sport. British Journal of Sports Medicine, 39(6), 319–323. https://doi.org/10.1136/bjsm.2005.018549

Piątek-Krzywicka, E., Borzucka, D., & Kuczyński, M. (2022). Postural control through force plate measurements in female AIS patients compared to their able-bodied peers. Scientific Reports, 12(1), 13170. https://doi.org/10.1038/s41598-022-17597-y

Raffalt, P. C., Fillingsnes Marker, I., Adler, A. T., & Alkjaer, T. (2021). Dynamics of Postural Control in Elite Sport Rifle Shooters. Journal of Motor Behavior, 53(1), 20–29. https://doi.org/10.1080/00222895.2020.1723478

Richer, N., & Lajoie, Y. (2020). Automaticity of Postural Control while Dual-tasking Revealed in Young and Older Adults. Experimental Aging Research, 46(1), 1–21. https://doi.org/10.1080/0361073X.2019.1693044

Roemmich, R. T., & Bastian, A. J. (2022). Motor control: In constant pursuit of optimality. In Current Biology (Vol. 32, Issue 10, pp. R462–R463). https://doi.org/10.1016/j.cub.2022.03.058

Sannicandro, I., Cofano, G., Rosa, R. A., & Piccinno, A. (2014). Balance training exercises decrease lower-limb strength asymmetry in young tennis players. Journal of Sports Science & Medicine, 13(2), 397–402. http://www.ncbi.nlm.nih.gov/pubmed/24790496

Saraiva, M., Vilas-Boas, J. P., Fernandes, O. J., & Castro, M. A. (2023). Effects of Motor Task Difficulty on Postural Control Complexity during Dual Tasks in Young Adults: A Nonlinear Approach. Sensors, 23(2). https://doi.org/10.3390/s23020628

Sipko, T., & Kuczyński, M. (2013). Intensity of chronic pain modifies postural control in low back patients. European Journal of Pain (United Kingdom), 17(4), 612–620. https://doi.org/10.1002/j.1532-2149.2012.00226.x

Söderman, K., Werner, S., Pietilä, T., Engström, B., & Alfredson, H. (2000). Balance board training: prevention of traumatic injuries of the lower extremities in female soccer players? A prospective randomized intervention study. Knee Surgery, Sports Traumatology, Arthroscopy : Official Journal of the ESSKA, 8(6), 356–363. https://doi.org/10.1007/s001670000147

Srihi, S., Jouira, G., Ben Waer, F., Rebai, H., Majdoub, A., & Sahli, S. (2022). Postural Balance in Young Tennis Players of Varied Competition Levels. Perceptual and Motor Skills, 129(5), 1599–1613. https://doi.org/10.1177/00315125221108913

Strang, A. J., Haworth, J., Hieronymus, M., Walsh, M., & Smart, L. J. (2011). Structural changes in postural sway lend insight into effects of balance training, vision, and support surface on postural control in a healthy population. European Journal of Applied Physiology, 111(7), 1485–1495. https://doi.org/10.1007/s00421-010-1770-6

Szafraniec, R., Barańska, J., & Kuczyński, M. (2018). Acute effects of core stability exercises on balance control. Acta of Bioengineering and Biomechanics, 20(4), 145–151. http://www.ncbi.nlm.nih.gov/pubmed/30520448

Terada, M., Kosik, K., Johnson, N., & Gribble, P. (2018). Altered postural control variability in older-aged individuals with a history of lateral ankle sprain. Gait and Posture, 60, 88–92. https://doi.org/10.1016/j.gaitpost.2017.11.009

Verhagen, E. A. L. M., Van der Beek, A. J., Bouter, L. M., Bahr, R. M., & Van Mechelen, W. (2004). A one season prospective cohort study of volleyball injuries. British Journal of Sports Medicine, 38(4), 477–481. https://doi.org/10.1136/bjsm.2003.005785

Watanabe, T., Nojima, I., Sugiura, H., Yacoubi, B., & Christou, E. A. (2019). Voluntary control of forward leaning posture relates to low-frequency neural inputs to the medial gastrocnemius muscle. Gait & Posture, 68, 187–192. https://doi.org/10.1016/j.gaitpost.2018.11.026

Winter, D. A., Patla, A. E., Prince, F., Ishac, M., & Gielo-Perczak, K. (1998). Stiffness control of balance in quiet standing. Journal of Neurophysiology, 80(3), 1211–1221. https://doi.org/10.1152/jn.1998.80.3.1211

Zemková, E., & Kováčiková, Z. (2022). Sport-specific training induced adaptations in postural control and their relationship with athletic performance. Frontiers in Human Neuroscience, 16, 1007804. https://doi.org/10.3389/fnhum.2022.1007804

